# Humanized APOE mouse brain volume increases over age irrespective of sex and APOE genotype: Implications for translational validity to the human

**DOI:** 10.1101/2025.10.27.684892

**Authors:** Adam C. Raikes, Avnish Bhattrai, Tian Wang, Jean-Paul Wiegand, Roberta Diaz Brinton

**Affiliations:** Center for Innovation in Brain Science, University of Arizona, Tucson, AZ, United States; Department of Neurology, University of Arizona, Tucson, AZ, United States; Department of Pharmacology, College of Medicine Tucson, University of Arizona, Tucson, AZ, United States

## Abstract

Translational validity of mouse models of human aging and late-onset Alzheimer’s disease (LOAD) risk are essential for both fundamental mechanistic science and therapeutic development. Given that the strongest risk factors for LOAD are age, female sex, and APOE-ε4 carriership, models must reflect these biological variables and disease phenotypes. The use of mouse models with humanized APOE (hAPOE) is a key strategy to advance translational validity.

To initially address translational validity of the hAPOE mouse model, we conducted ex-vivo magnetic resonance imaging analysis of brain volumes in male and female mice across APOE genotypes (ε3/ε3, ε3/ε4, ε4/ε4) and ages corresponding to a human lifespan of approximately 30-70 years (6-25 months in mice). The primary outcomes indicated that total MRI brain volume increased with age (an average of 2.12mm3 per month), irrespective of sex or *APOE* genotype. Additionally, APOE-ε4 carriers had greater total brain volumes than non-carriers. No sex differences were observed in total brain volume.

Voxelwise analyses revealed a pattern of localized morphometric changes independent of differences in total brain volume. Age-related volumetric increases were distributed across subcortical regions (e.g., thalami, hippocampi), while age-related decreases were evident across cortical regions, notably the bilateral parieto-temporal and frontal lobes. Additionally, sex differences were evident after controlling for total brain volume, with females showing greater cortex-dominant volumes while males showed a pattern of greater volumes in regions including cerebellar cortices, olfactory bulbs, and striata. No genotypic effects were observed in the voxelwise analysis after correcting for multiple comparisons, suggesting that *APOE* genotype does not drive localized volume differences independent of total brain volume.

These findings indicate that, at the MRI level of analysis, this humanized *APOE* mouse model does not recapitulate the volumetric atrophy typically seen in human brain aging and Alzheimer’s disease. The results suggest that humanized *APOE* alone is insufficient to induce the LOAD atrophy phenotype. This model may better serve as a platform for studying vulnerable aging rather than a primary model for progressive neurodegenerative atrophy.

## Introduction

Approximately 1 in 9 Americans over the age of 65 lives with late-onset Alzheimer’s disease (LOAD)^1^. Without effective treatments, the prevalence is expected to exceed 12.7 million by 2050 with new diagnoses exceeding 1 million per year^1–3^. The strongest risk factors for LOAD are age, female sex, and APOE-ε4 carriership^4–6^. Therapeutic development requires disease models that reflect the biological risk factors as well as disease-like phenotypes. As noted in two recent reviews, the majority of mouse models used are the 5xFAD and the 3xTg, which represent the disease process well in terms of amyloid deposition and atrophy, but are suboptimal in terms of their representativeness of human disease^7,8^. Despite the fact that these mice replicate the disease process, the individual genetic mutations required are rare and not observed in combination in the human population^9,10^. Single mutations result in an early age of disease onset (translating to approximately 30 years of age in the human, mimicking early-onset AD, rather than late-onset), an inconsistent disease profile in male mice, and mouse Apoe rather than human APOE, which does not confer the same disease mechanisms^7–9,11^. Given that the APOE-ε4 allele is the strongest genetic risk factor for LOAD, rodent models that include humanized APOE are essential for translationally relevant therapeutic development.

To date there are multiple targeted replacement, knock-out, and knock-in models for APOE mice. Although incorporating a major genetic risk factor for AD, these mice do recapitulate key aspects of disease, including cognitive impairment^12,13^. To address the paucity of AD relevant pathology, multiple APOE models have been developed that include other non-dominant mutations, including *Trem2*^14,15^, humanized *NOS*^16,17^, or humanized *APP*^18^. Although beneficial for driving pathology, the isolated role of APOE in AD risk and progression is not clearly identifiable in these models.

To this end, a knock-in mouse model expressing humanized APOE-ε3/ε3 and APOE-ε4/ε4 has been developed by the MODEL-AD Consortium (model-ad.org). Recent work with this model demonstrates that the APOE-ε4/ε4 mice have reduced cerebral metabolism and neurovascular uncoupling^19^, altered peripheral metabolism^20,21^, immunologic profiles^12,20,22^, and transcriptomic profiles^21,23^, decreased cortical microglial responsiveness^24^, accelerated endocrinological aging^25^, as well as subtle cognitive differences^12,13^. Many of these effects are sex-specific^21–23,25^, which is consistent with differences in the sex distribution of Alzheimer’s disease. These findings suggest that, in some respects, this mouse model may be a suitable risk model for Alzheimer’s disease.

To date, there are no studies that deeply characterize MRI brain imaging phenotypes of aged humanized APOE-ε3/ε3 and APOE-ε4/ε4. Past work with mouse models that include humanized APOE in addition to other disease- and inflammation-promoting genetic profiles has demonstrated volumetric decreases associated with the APOE-ε4 allele^14,16,26,27^ as well as decreased fractional anisotropy in major white matter tracts, the piriform, and hippocampus^14,16^. However, the extent of these differences is not consistently reported. For example, Badea et al (2022) and Badea et al (2019) reported no decreases in hippocampal volume, while Yin et al (2014) reported lower volume for APOE-ε4 mice^16,26,27^. These contradictory findings are similar to those reported for APOE-ε4 FAD models^8^ as well as 3xTg and 5xFAD models^7^. Important confounding effects should be noted for these studies, including other genetic mutations, inconsistent control groups, and varied methods for adjusting regional volumes for total brain volume. Collectively, these factors limit the interpretability of APOE-related effects in these mouse models as well as the translational potential for therapeutic development.

The purpose of the present study was to investigate the effects of age, sex, and APOE genotype on total brain and localized brain volumes in the MODEL-AD humanized APOE mouse model. As a candidate AD risk model, we hypothesized that humanized APOE-ε4/ε4 mice would exhibit evidence of atrophy and that this effect would be exacerbated in female mice compared to males.

## Materials and Methods

### Animals

All animal studies were performed following National Institutes of Health guidelines on the use of laboratory animals and all protocols were approved by the University of Arizona Institutional Animal Care and Use Committee. Homozygous humanized APOE-ε4 targeted replacement mice were obtained from Jackson Laboratory (#027894). Humanized APOE-ε3 target replacement heterozygous mice were obtained from Jackson Laboratory (#029018) and bred to get homozygous APOE-ε3 mice. Mice were housed in sterile conditions on 14-hour light, 10-hour dark cycles and provided *ad libitum* access to food and water.

### Image acquisition

Mice were sacrificed at prespecified ages (6, 9, 15, and 23–25 months) for ex vivo MRI. Each animal underwent transcardial perfusion with phosphate-buffered saline (PBS) for 4–6 minutes, followed by perfusion with 4% paraformaldehyde (PFA) in PBS for an additional 4–6 minutes. After perfusion, skulls were cleaned of soft tissue and stored in 4% PFA in PBS for 48 hours to ensure complete fixation and enable in-skull imaging. Samples were then transferred to 0.01% sodium azide for long-term storage. Prior to MRI, fixed skulls were placed in vacuum-desiccated 10 mL syringes filled with Fluorinert (FC-3283) for imaging. High resolution structural images were acquired with a T2-weighted rapid acquisition with relaxation enhancement (RARE) sequence (TE: 30 ms; TR: 1800 ms; flip angle 180°; RARE factor: 8; number of averages: 2; FOV: 2.4 × 1.44 × 0.96; acquisition matrix 320×192×120; reconstructed voxel size 75μm; total acquisition time: 3 hours and 4 minutes).

### Image minimal pre-processing

All preprocessing on and analyses were conducted on the high performance computing cluster at the University of Arizona. All data were preprocessed using a custom pipeline written in Bash and Python. A custom Apptainer container was created with necessary softwares for version control and was used unless otherwise noted. This included ANTs^28^ (v. 2.4.4), FSL^29^ (v. 6.0.6.4), MRtrix^30^ (v. 3.0.4), minc-toolkit-extras (https://github.com/CoBrALab/minc-toolkit-extras; commit 544485d), *optimized_antsMultivariateTemplateConstruction* (https://github.com/CoBrALab/optimized_antsMultivariateTemplateConstruction; commit ca4ba15) and Python (v 3.10.12).

### Anatomical preprocessing

T2-weighted images underwent initial automated quality control^31^. Outlier images were examined visually and images with significant non-uninform intensities (n = 4) were excluded. All included images were reoriented to RAS ordering and intensity-normalized using percentile-based histogram rescaling. Foreground estimation was performed via Otsu thresholding, followed by centering, padding, and cropping to a constrained field of view. Bias field correction was applied using N4BiasFieldCorrection^32^, with masks refined through iterative affine and nonlinear registration to the DSURQE^33–36^ mouse brain template (https://github.com/IBT-FMI/mouse-brain-templates) resampled to 0.075mm isotropic voxel size. Non-local means denoising^37^ preceded final bias correction and intensity rescaling. Brain masks were generated by inverse warping of the template mask, yielding a final preprocessed T2w image and corresponding brain mask.

### Deformation based analysis

An initial minimum deformation template was built using *optimized_antsMultivariateTemplateConstruction*. A total of 96 mice were used to build the template and included 4 mice from each age (6, 9, 15, 24 months), genotype (hAPOE-ε3/3, hAPOE-ε3/4, hAPOE-ε4/4), and sex (male, female) combination. The process began by building an initial template using six iterations of rigid-only transformations to align the images. The resulting average from the final rigid iteration was then used as the target for the next stage, which built a new template using six iterations of similarity transformations. This procedure was sequentially repeated for six iterations of affine transformations and finally for six iterations of non-linear Symmetric Normalization (SyN) transformations. At each iteration of every stage, all subjects were registered to the target template; a new template was then generated by averaging the warped images, sharpening the result, and applying the scaled, inverted average of the transformations for use in the next iteration. A final consensus mask was also generated, computed as the mean of the template-space transformed brain masks and thresholded to include voxels in at least 50% of the masks.

After this process, all of the brains (n = 159) were independently re-registered to the minimum deformation template using a single iteration step identical to the template building process, which included both linear (rigid, affine) and nonlinear (SyN) registration components.

Deformation based morphometry was implemented through *optimized_antsMultivariateTemplateConstruction* to produce absolute and relative log-transformed Jacobian determinants of the deformation fields transforming each individual mouse brain into the study-specific template space. Log-transformed Jacobians describe the amount of deformation to warp the template voxel to the individual mouse such that values > 0 indicate larger individual volumes and values < 0 indicate larger template volumes. The relative Jacobians, which exclude the affine and residual affine components of the transformations, were used for all analyses and intrinsically control for total brain size. All images were smoothed with a 4 voxel full width at half maximum kernel prior to analyses.

### Statistical analyses

Primary analyses were conducted in template voxel space using Convoxel and ModelArray^38,39^ (v. 0.1.5). ModelArray runs in R^40^ (v. 4.1.2). Voxelwise linear additive models were fit using ModelArray to examine the independent effects of age, sex, and genotype with sum to zero contrast coding for interpretable coefficients. Specifically, age was Z-scored (mean = 13.4 ± 5.94 months), sex compared males - females, and genotypic effects compared hAPOE-ε3/4 - hAPOE-ε4/4 and then hAPOE-ε3/3 - hAPOE-ε4 carriers. Voxelwise statistical significance was set at a two-sided *p*_*FDR*_ < 0.05. Statistically significant voxels were separated into positive and negative directionality (depending on the contrast; i.e. positive: male > female, negative: female > male), localized using the Dorr mouse brain atlas which had been transformed to study-specific template space and summarized as the percent of the region occupied by significant voxels (regional specificity) and the percent of the significant voxels represented by the region (pattern representation). Total brain volumes were additionally analyzed by inverse warping the template space mask to the individual mouse brains, summing the voxels and fitting a linear model with the same covariates as the voxelwise models. Plots were generated using a combination of software including *ggprism*^41^ (Figure 1), *nilearn*^42^ (Figure 2), and *tidyplots*^43^ (Figures 3-4).

**Figure 1.**
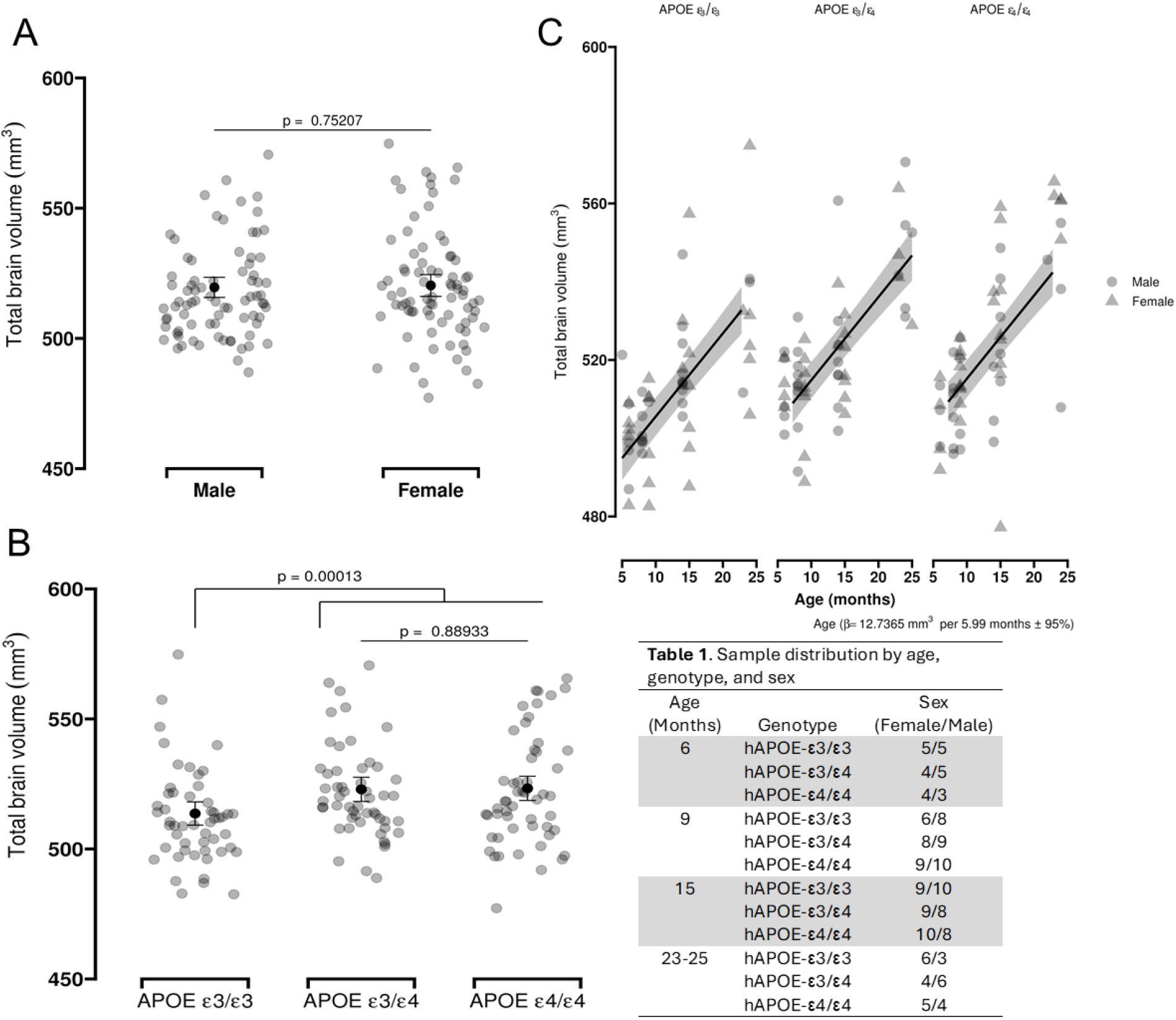
Sex, genotype, and age effects on total brain volume. A) There were no statistically significant differences between male and female mice. B) APOE-ε4 carriers exhibited greater volume than non-carriers. C) A statistically significant increase in brain volume as observed with increasing age. Group mean and error bars are model-derived marginal means + 95% CI (A,B). Line and ribbon (C) represents model fit + 95% CI.

**Figure 2.**
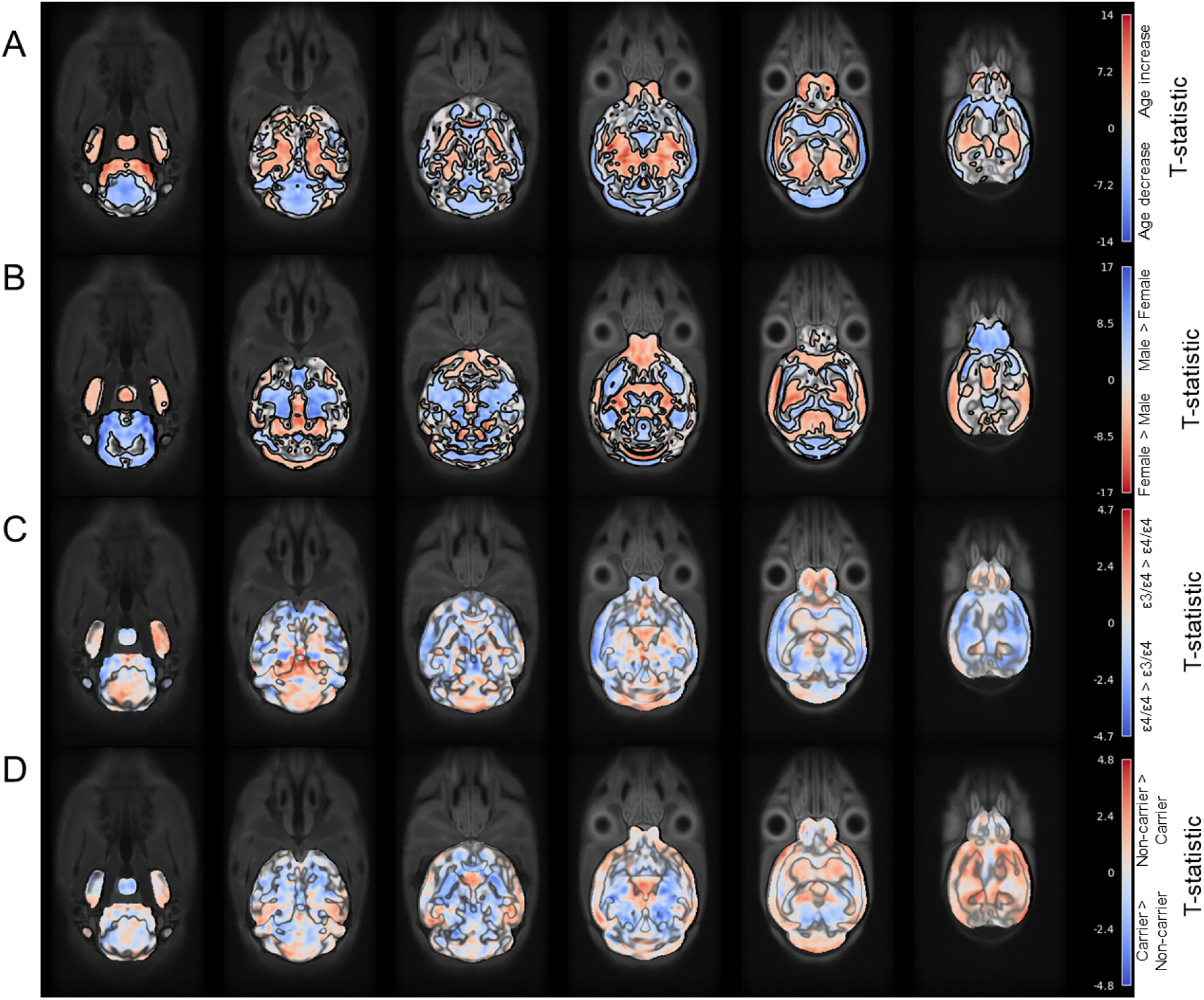
Voxelwise t-statistic maps for main effects of age (A), sex (B), APOE-ε4 hetero-vs. homozygosity (C), and APOE-ε4 carriership (D). Statistically significant findings (pFDR < 0.05) are outlined and opaque for age (A) and sex (B). Genotypic effects (C, D) did not survive multiple comparisons correction.

**Figure 3.**
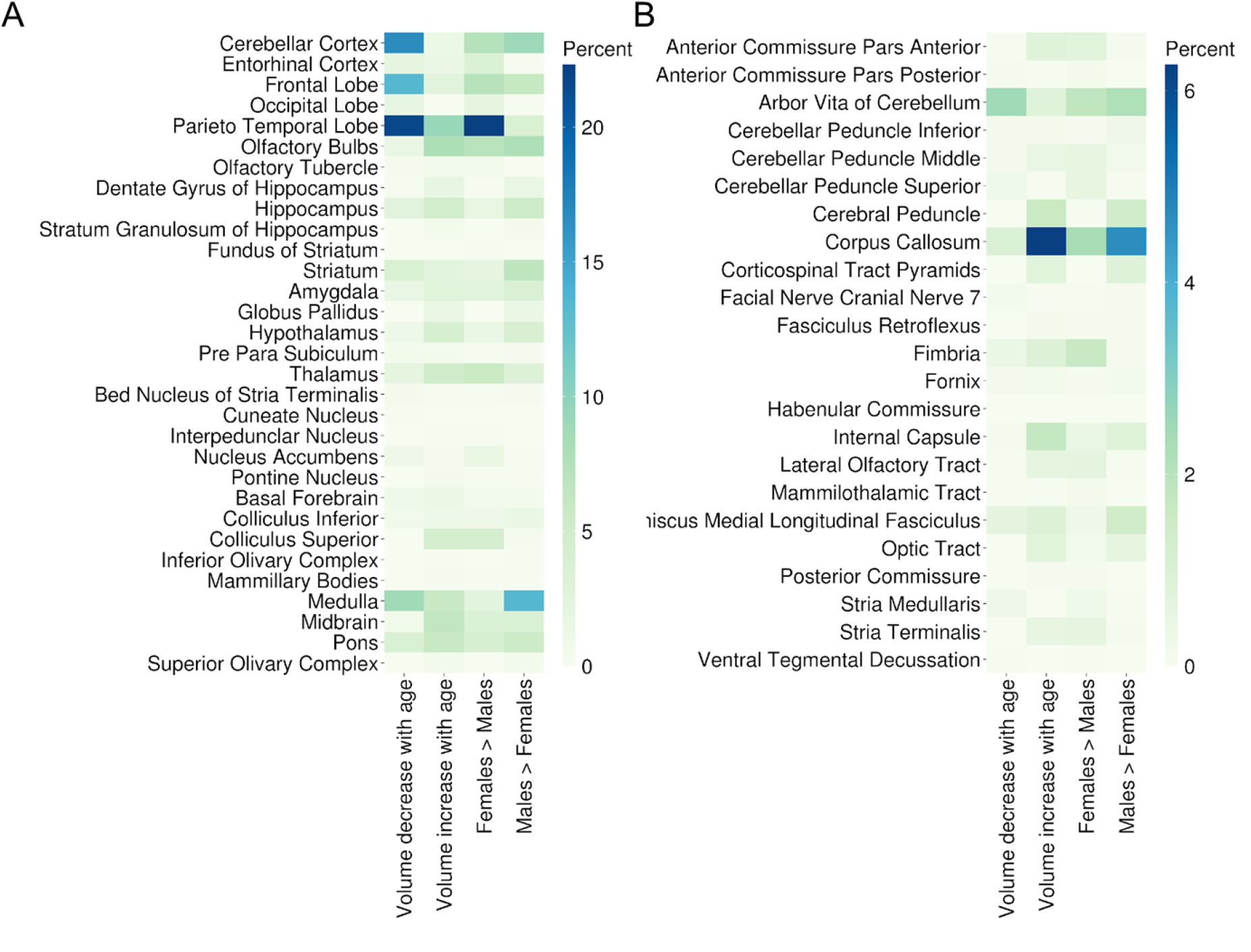
Pattern representation in gray (A) and white (b) matter. For each effect (age decrease, increase; sex differences), the heatmap indicates the proportion of statistically significant voxels within each region (i.e. >20% of the voxels in demonstrating an age-related decrease in volume were located in the the parieto-temporal lobe).

**Figure 4.**
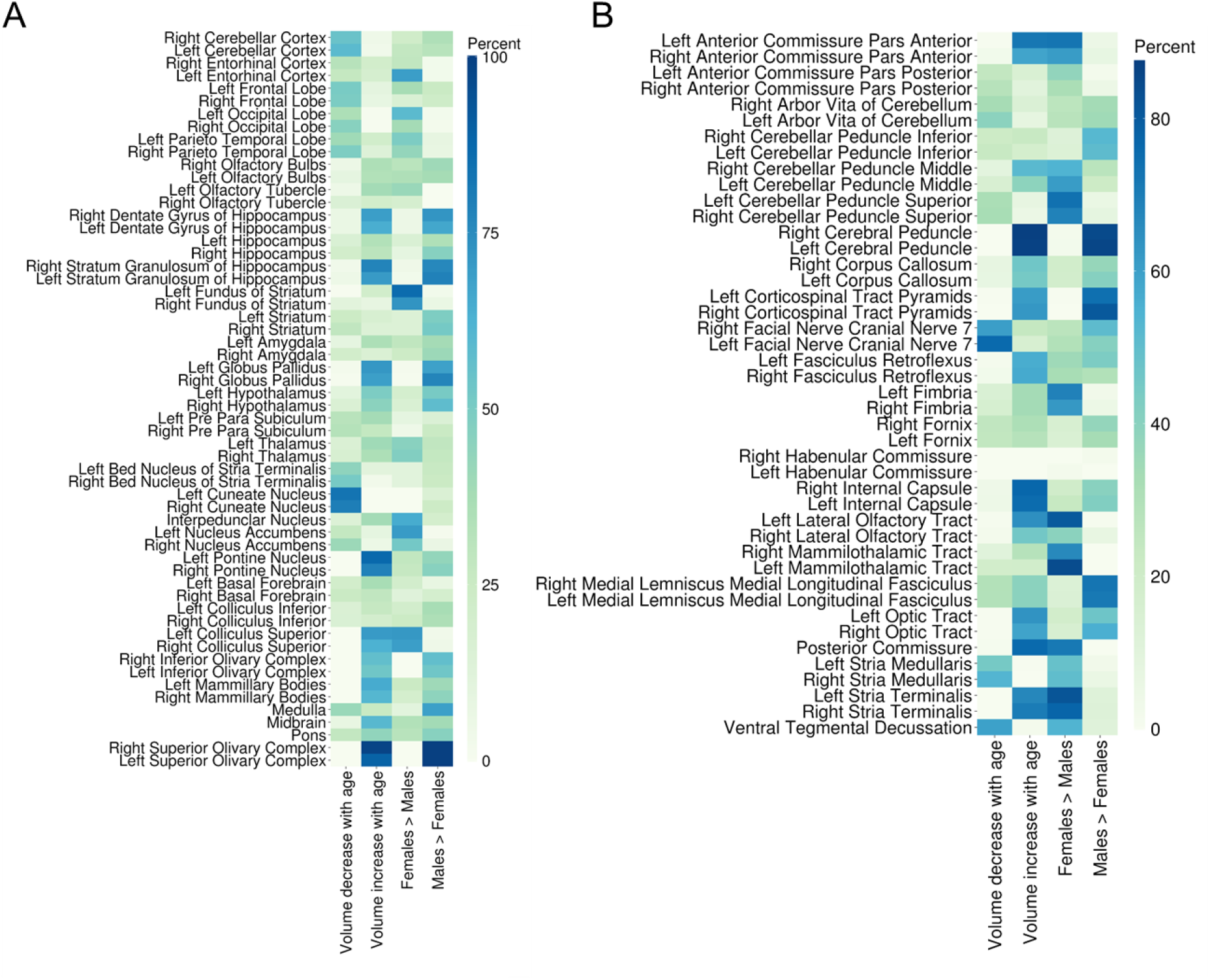
Regional specificity of effects for gray (A) and white (b) matter. For each effect (age decrease, increase; sex differences), the heatmap indicates the proportion of the regions voxels that are statistically significant (i.e., >80% of the voxels in the bilateral cerebral peduncles were larger in males compared to females.

## Results

### Cohort characteristics

A total of 158 mice were included in the present analyses. Distributions by age, sex, and genotype are presented in Figure 1. The linear model for the total brain volumes demonstrated statistically significant increases in brain volumes over the range of ages (6-25 months), with an average increase of 2.12mm^3^ per month. After controlling for all other parameters, APOE-ε4 carriers had greater total brain volumes than non-carriers (mean difference: 9.78mm^3^). No statistically significant differences were observed in total brain volumes between APOE-ε3/4 and APOE-ε4/4 mice or between males and females. Total brain volume analyses are summarized in Figure 1.

### Aging is associated with cortical decreases and subcortical preservation

Age-specific effects are presented in Figure 2A. Age-related increases were primarily observed in subcortical regions – including corpus callosum, thalami, hippocampi, superior colliculi, hypothalami, and amydalae – as well as a portion of the bilateral parieto-temporal lobe accounting for approximately 40% of the total number of significant voxels. Notably while this distribution of age-positive voxels was observed across multiple regions, the regional specificity (proportion of a region with significant voxels) was relatively smaller. Smaller gray matter nuclei (e.g., superior olivary complex, pontine nuclei) as well as smaller white matter pathways (anterior commissure, stria terminalis) and gray matter substructures (dentate and stratum granulosum of the hippocampus) demonstrated age-related regional specificity, with significant voxels in > 60% of those individual regions. Figures 3 and Figure 4 provide comparative heatmaps representing regional pattern representation and specificity.

The age-related pattern of decreasing volume was substantially represented by the bilateral parieto-temporal cortex (21.7% of significant voxels), cerebellar cortex (16.66% of significant voxels), and frontal lobes (13.39% of significant voxels). To a lesser extent, other regions, including the bilateral striata, hippocampi, entorhinal cortex, amygdala, and thalami were also represented in the overall pattern. Notably, these cortical and cerebellar cortical regions, as well as the occipital lobe, also demonstrated high regional specificity, with up to 50% of the regional voxels demonstrating age-related decrease. Figure 3 and Figure 4 provide comparative heatmaps representing regional pattern representation and specificity.

### Cortical and subcortical structures exhibit sexual dimorphism

Sex-specific effects are presented in Figure 2B. The regional pattern of male positive effects included the bilateral cerebellar cortices, olfactory bulbs, striata, frontal cortex, hippocampi, and corpus callosum. High regional specificity was observed in the bilateral olivary complexes, white matter pathways (cerebral peduncles, corticospinal tract pyramids, medial longitudinal fasciculus) as well as subcortical structures (globus pallidus, bilateral stratum granulosa and dentate gyri of the hippocampus). Figure 3 and Figure 4 provide comparative heatmaps representing regional pttern representation and specificity.

In contrast, females exhibited greater cortex-dominant regional patterns, with bilateral parieto-temporal, frontal, cerebellar, and entorhinal cortices all substantially contributing to the overall female-specific pattern. White matter tracts (mammilothalamic, stria terminalis, olfactory, cerebellar peduncle superior, anterior commissure), bilateral subependymale zones rhinocele, and the entorhinal cortex further contributed to this female-specific pattern. Figure 3 and Figure 4 provide comparative heatmaps representing regional pattern representation and specificity.

### APOE genotype does not drive localized volumetric differences

No voxelwise differences survived multiple comparisons correction for any of the APOE genotype analyses, after accounting for age and sex effects (Figure 2C-D).

## Discussion

### Key findings

The purpose of this work was to determine the translational relative of MRI-based volumetric profiles of the humanized APOE male and female brain. This mouse model was developed as a candidate for translationally relevant research in Alzheimer’s disease addressing the hypothesis that APOE genotype would be a significant driver of Alzheimer’s-like morphological phenotypes. Brain MRI imaging findings reported herein indicate A) that age and APOE-ε4 carriership significantly increase total brain volumes in this mouse model; B) that localized age-related changes in brain morphometry are independent of the global age-related increase, with widespread decreases particularly in cortical volume; and C) that sexually dimorphic patterns of local volume differences exist. However, contrary to expectations, APOE genotype did not impact regional volumes in the same manner observed in human studies.

### Total brain volume

The present findings indicate that brain volumes in this mouse model continue to increase over the lifespan. This is consistent with findings in different mouse models, including the background C57/B6 strain, where rapid brain growth is observed over the first few months of life followed by a much slower increase or plateauing of brain volumes over the course of aging^7^. These findings are dissimilar to findings in human aging, where the brain reaches peak volume within the first two decades of life, followed by plateauing or decline in volume over the next 50-70 years^44^. Normal whole brain volume may decline at rates of up to < 0.5% per year, depending on decade of life^45^ and requires between 30 and 35 years for volume to decrease by two standard deviations from its value at age 40 during healthy aging^46^. In AD, the rate of decline is accelerated^44^. The cross-sectional nature of our mouse data does not permit an estimate of rates of growth or atrophy, but the observed increase in volumes across the 18 months represented here indicates an estimated 7% increase in total volume. Notably, the brain mask for estimating total volume includes the ventricles, which may contribute to this increase, whereas ex-vivo imaging here does not permit accurate evaluation of ventricular size^47^.

Independent of the observed aging effects, a small, statistically significant amount of the variability in total brain volume was explained by genotypic variation, with APOE-ε4 carriers having greater brain volumes than non-carriers. The observed magnitude (9.72mm^3^) is small and may not represent a translationally relevant magnitude of difference. For reference, this magnitude of difference is approximately the same size as the left thalamus distributed over the entire mouse brain. While APOE-ε4 mediated whole brain atrophy is evident in Alzheimer’s disease, recent studies using large cohorts reflecting presumably healthy aging have revealed subtle or absent APOE genotypic effects on total gray matter and total brain volume^48–50^, which is broadly consistent with the findings here.

Interestingly, after accounting for age and APOE genotype, no sex differences were evident in total brain volumes. Sex differences in total brain volume are robustly reported across the human literature, yet absent in the present analyses^44,51–53^. MRI findings from the background strain (C57BL6) show the same lack of total brain volume differences as observed here^54^, suggesting that this is not a by-product of the inclusion of humanized APOE but rather a mouse-specific phenotype. Collectively, these results indicate that this mouse model may better represent aging processes at the whole brain level rather than genetic trait related phenotypes.

### Aging results in decreased cortical and preserved or increased subcortical volume

While overall increases in brain volume occurred over the aging process, local changes in volume occurred independently of this increase. Specifically, the cortex exhibited decreased volume while subcortical regions exhibited more frequently preserved or increased volumes. This age-related pattern is evident in both human^44,55^ and mouse^56^ literature. Prior work in a related but different mouse model suggests the presence of both metabolic deficits and altered neurovascular coupling in the cortex, which may provide a plausible explanation for the decreased cortical volume with advancing age^15,19^. Notably, the present mouse model does not display any of the hallmark pathological features of neurodegenerative processes, and thus cortical-subcortical shift may better reflect non-pathological aging rather than neurodegenerative atrophy. However, the localization and regional specificity of some of the volumetric decline – particularly the parieto-temporal and entorhinal cortices – does reflect early atrophy sites in Alzheimer’s disease^57–61^. This specific decline may, therefore, reflect some AD-like vulnerability, rather than pathology.

### Cortical and subcortical structures exhibit sexual dimorphism

Interestingly, sex differences were observed in local volumes. Females generally had greater cortical volumes, independent of any age-related decline, than males. These findings are consistent with the pattern of sex differences observed in a comparative study of humans and C57BL6 mice^54^, in which consistent sex differences were observed across species with subcortical and deep gray brain structures exhibiting a male bias where cortical regions exhibited a female bias. These findings are generally consistent with other human^51,62^ and mouse studies^63^ of sex differences. These sex-biased regional variations may likely reflect sex hormone exposure and timing in both species^63,64^.

### APOE genotype does not drive local volume variation

Contrary to expectation, APOE genotype did not exert significant voxel-level effects. Analyses herein controlled for the total brain volume indicating that total brain volume differences were more global in nature rather than driven by regionally focal genotypic variation.

Prior work with this specific mouse model has revealed APOE-ε4/ε4 decreases in cerebral metabolism with aging^19^, alterations in neuronal^65^ and peripheral metabolism^20,21^, immunologic profiles^12,20,22^, cortical transcriptomic profiles^21,23^, heterogeneous weight trajectories across aging^66^, and accelerated endocrine aging^25^. Mutations (*Trem*2^15^; *Trem2* + *hAPP*^67^) in addition to humanized APOE coupled with environmental factors (high fat diet)^67^ revealed reduced cerebral glucose metabolism and neurovascular coupling. In hAPOE-ε4s carrying both *Trem2* and *hAPP* mutations, a high fat diet induced volumetric decreases compared to a normal control diet at multiple age points^67^. Given those findings, results reported herein indicate that humanized APOE-ε4 alone is insufficient to induce Alzheimer’s-like MRI phenotypes of atrophy.

Collectively, and in the absence of discernible pathology, these findings point to an aging process characterized by widespread cortical volumetric reductions that are coincident with increased subcortical volumes. These changes are further modified by sex, with female preservation of cortical volumes and male preservation of subcortical volumes, without any specific impact of humanized APOE genotype. These findings partially echo findings from more aggressive AD mouse models that do exhibit pathological hallmarks (amyloid B deposition, tau)^7,8^. In those models, age-related cortical thinning and volumetric decrease are greater and more accelerated compared to control mice^7,8^. However, the present humanized APOE model, without other aggressive disease-causing mutations, fails to recapitulate the extent of progressive atrophy observed in aggressive, but translationally constrained, mouse models. Rather, these mice appear to represent a model of vulnerable aging, particularly with APOE-ε4/ε4 females demonstrating accelerated aging^25^, and may serve as a reasonable aging control group when evaluating more aggressive disease models.

### Strengths and limitations

This study capitalizes on a number of strengths. Here we pool from a large number of humanized APOE mice across an extensive age range, reflecting an estimated human age range of 30-70. Our imaging pipeline capitalizes on open-source softwares to enable high-throughput processing, robust brain masking and template registration, as well as voxel-wise analyses and ultimately reproducibility within and across research groups. The template brains built from this analytic process can serve as a basis for future target registrations and comparative analyses with other mouse models. However, several limitations should be acknowledged. First, the humanized APOE-ε3/3 mouse was our control group, rather than a wild-type mouse. While the inclusion of wild type mice may have revealed greater hAPOE effects, the comparison would then have been between mouse-type Apoe and human APOE and we did not view this as a clinically or translationally relevant model. Second, we did not perform behavioral phenotyping in all of the mouse cohorts, thus we could not investigate whether these brain morphometric differences were related to relevant behavioral outcomes. Third, all animals were housed in sterile environments with a controlled, standard diet. This housing strategy decreases the potential influence of environmental confounders (e.g., bacterial and viral contagions, pollution) on the genetic risk factors at the sacrifice of real world environmental factors that may contribute to disease onset and progression in humans^9,68^. Finally, nuclear (i.e. FDG-PET) and molecular (i.e. IHC) assays were not available for the full cohort of animals. Therefore, mechanistically linking macroscale changes in MRI features to more granular causal factors was not feasible.

## Conclusions

Volumetric analyses of the humanized APOE mouse model indicate that these mice do not present with an Alzheimer’s disease-like phenotype, despite disease-related risk factors of age, female sex, and APOE genotype. These findings suggest that, for these mice, APOE-ε4 alone does not confer macrostructural atrophy and likely requires additional risk factors (genetic and/or environmental). Analyses are underway with a mouse model that combines humanized APOE genotypes with humanized APP, which may better model the amyloid-generating process and ultimately disease progression.

## Acknowledgements

We appreciate Dr. Beth Hutchinson, Dr. Ted Trouard, and Dr. JP Galons for their expertise in setting and optimizing our MRI acquisition protocols and for their input on preprocessing and analyses.

## Funding

This work was supported by the NIH/National Institute on Aging grants P01AG026572 and R01AG057931 to RDB and the Center for Innovation in Brain Science to RDB. Neuroimaging was supported by the University of Arizona’s Translational Bioimaging Resource (TBIR) and NIH small instrumentation grant (S10 OD025016). Neuroimaging preprocessing and statistical analyses were supported by High Performance Computing (HPC) resources supported by the University of Arizona TRIF, UITS, and Research, Innovation, and Impact (RII) and maintained by the UArizona Research Technologies department.

## Notes

### Competing Interest Statement

The authors have declared no competing interest.

## References

1. 2025 Alzheimer’s disease facts and figures. Alzheimers Dement. 2025;21(4):e70235. doi:10.1002/alz.70235

2. CDC. About Alzheimer’s. Alzheimer’s Disease and Dementia. February 18, 2025. Accessed October 15, 2025. https://www.cdc.gov/alzheimers-dementia/about/alzheimers.html

3. Fang M, Hu J, Weiss J, et al. Lifetime risk and projected burden of dementia. Nat Med. 2025;31(3):772–776. doi:10.1038/s41591-024-03340-9

4. Riedel BC, Thompson PM, Brinton RD. Age, APOE and sex: Triad of risk of Alzheimer’s disease. J Steroid Biochem Mol Biol. 2016;160:134–147. doi:10.1016/j.jsbmb.2016.03.012

5. O’Neal MA. Women and the risk of Alzheimer’s disease. Front Glob Womens Health. 2024;4. doi:10.3389/fgwh.2023.1324522

6. Liu CC, Kanekiyo T, Xu H, Bu G. Apolipoprotein E and Alzheimer disease: risk, mechanisms and therapy. Nat Rev Neurol. 2013;9(2):106–118. doi:10.1038/nrneurol.2012.263

7. Jullienne A, Trinh MV, Obenaus A. Neuroimaging of mouse models of alzheimer’s disease. Biomedicines. 2022;10(2):305. doi:10.3390/biomedicines10020305

8. Balu D, Karstens AJ, Loukenas E, et al. The role of APOE in transgenic mouse models of AD. Neurosci Lett. 2019;707:134285. doi:10.1016/j.neulet.2019.134285

9. Granzotto A, Vissel B, Sensi SL. Lost in translation: inconvenient truths on the utility of mouse models in alzheimer’s disease research. Behrens TE, ed. Elife. 2024;13:e90633. doi:10.7554/eLife.90633

10. Tanzi RE. The Genetics of Alzheimer Disease. Cold Spring Harb Perspect Med. 2012;2(10):a006296. doi:10.1101/cshperspect.a006296

11. Tai LM, Balu D, Avila-Munoz E, et al. EFAD transgenic mice as a human APOE relevant preclinical model of Alzheimer’s disease. J Lipid Res. 2017;58(9):1733–1755. doi:10.1194/jlr.R076315

12. Sepulveda J, Luo N, Nelson M, Ng CAS, Rebeck GW. Independent APOE4 knock-in mouse models display reduced brain APOE protein, altered neuroinflammation, and simplification of dendritic spines. J Neurochem. 2022;163(3):247–259. doi:10.1111/jnc.15665

13. McLean JW, Bhattrai A, Vitali F, Raikes AC, Wiegand JPL, Brinton RD. Contributions of sex and genotype to exploratory behavior differences in an aged humanized APOE mouse model of late-onset Alzheimer’s disease. Learn Mem. 2022;29(9):321–331. doi:10.1101/lm.053588.122

14. Shang Y, Mishra A, Wang T, et al. Evidence in support of chromosomal sex influencing plasma based metabolome vs APOE genotype influencing brain metabolome profile in humanized APOE male and female mice. Reddy H, ed. PLOS One. 2020;15(1):e0225392. doi:10.1371/journal.pone.0225392

15. Kotredes KP, Oblak A, Pandey RS, et al. Uncovering disease mechanisms in a novel mouse model expressing humanized APOEε4 and Trem2*R47H. Front Aging Neurosci. 2021;13:735524. doi:10.3389/fnagi.2021.735524

16. Badea A, Wu W, Shuff J, et al. Identifying Vulnerable Brain Networks in Mouse Models of Genetic Risk Factors for Late Onset Alzheimer’s Disease. Front Neuroinformatics. 2019;13:72. doi:10.3389/fninf.2019.00072

17. Badea A, Delpratt NA, Anderson RJ, et al. Multivariate MR biomarkers better predict cognitive dysfunction in mouse models of Alzheimer’s disease. Magn Reson Imaging. 2019;60:52–67. doi:10.1016/j.mri.2019.03.022

18. Badea A, Kane L, Anderson RJ, et al. The fornix provides multiple biomarkers to characterize circuit disruption in a mouse model of Alzheimer’s disease. Neuroimage. 2016;142:498–511. doi:10.1016/j.neuroimage.2016.08.014

19. Onos KD, Lin PB, Pandey RS, et al. Assessment of neurovascular uncoupling: APOE status is a key driver of early metabolic and vascular dysfunction. Alzheimers Dement. 2024;20(7):4951–4969. doi:10.1002/alz.13842

20. Fleeman RM, Snyder AM, Kuhn MK, et al. Predictive link between systemic metabolism and cytokine signatures in the brain of apolipoprotein E ε4 mice. Neurobiol Aging. 2023;123:154–169. doi:10.1016/j.neurobiolaging.2022.11.015

21. Foley KE, Diemler CA, Hewes AA, Garceau DT, Sasner M, Howell GR. APOE ε4 and exercise interact in a sex-specific manner to modulate dementia risk factors. Alzheimers Dement Transl Res Clin Interv. 2022;8(1):e12308. doi:10.1002/trc2.12308

22. Fleeman RM, Kuhn MK, Chan DC, Proctor EA. Apolipoprotein E ε4 modulates astrocyte neuronal support functions in the presence of amyloid-β. J Neurochem. 2023;165(4):536–549. doi:10.1111/jnc.15781

23. Foley KE, Hewes AA, Garceau DT, et al. The APOEε3/ε4 genotype drives distinct gene signatures in the cortex of young mice. Front Aging Neurosci. 2022;14:838436. doi:10.3389/fnagi.2022.838436

24. Sepulveda J, Kim JY, Binder J, Vicini S, Rebeck GW. APOE4 genotype and aging impair injury-induced microglial behavior in brain slices, including toward aβ, through P2RY12. Mol Neurodegener. 2024;19(1):24. doi:10.1186/s13024-024-00714-y

25. Wang T, Mao Z, Shang Y, et al. Accelerated midlife endocrine and bioenergetic brain aging in APOE4 females. Front Aging Neurosci. 2025;17:1632877. doi:10.3389/fnagi.2025.1632877

26. Badea A, Stout JA, Anderson RJ, Cofer GP, Duan LL, Vogelstein JT. Imaging Biomarkers for Alzheimer’s Disease Using Magnetic Resonance Microscopy. In: Haber-Pohlmeier S, Blümich B, Ciobanu L, eds. Magnetic Resonance Microscopy. 1st ed. Wiley; 2022:283–314. doi:10.1002/9783527827244.ch13

27. Yin J, Turner G, Coons S, Maalouf M, Reiman E, Shi J. Association of Amyloid Burden, Brain Atrophy and Memory Deficits in Aged Apolipoprotein ε4 Mice. Curr Alzheimer Res. 2014;11(3):283–290. doi:10.2174/156720501103140329220007

28. Tustison NJ, Cook PA, Holbrook AJ, et al. The ANTsX ecosystem for quantitative biological and medical imaging. Sci Rep. 2021;11(1):9068. doi:10.1038/s41598-021-87564-6

29. Jenkinson M, Beckmann CF, Behrens TEJ, Woolrich MW, Smith SM. FSL. Neuroimage. 2012;62(2):782–790. doi:10.1016/j.neuroimage.2011.09.015

30. Tournier JD, Smith R, Raffelt D, et al. MRtrix3: a fast, flexible and open software framework for medical image processing and visualisation. Neuroimage. 2019;202:116137. doi:10.1016/j.neuroimage.2019.116137

31. Kalantari A, Shahbazi M, Schneider M, et al. Automated quality control of small animal MR neuroimaging data. Imaging Neurosci. 2024;2:imag-2-317. doi:10.1162/imag_a_00317

32. Tustison NJ, Avants BB, Cook PA, et al. N4ITK: improved N3 bias correction. IEEE Trans Med Imaging. 2010;29(6):1310–1320. doi:10.1109/TMI.2010.2046908

33. Dorr AE, Lerch JP, Spring S, Kabani N, Henkelman RM. High resolution three-dimensional brain atlas using an average magnetic resonance image of 40 adult C57Bl/6J mice. Neuroimage. 2008;42(1):60–69. doi:10.1016/j.neuroimage.2008.03.037

34. Steadman PE, Ellegood J, Szulc KU, et al. Genetic effects on cerebellar structure across mouse models of autism using a magnetic resonance imaging atlas. Autism Res. 2014;7(1):124–137. doi:10.1002/aur.1344

35. Ullmann JFP, Watson C, Janke AL, Kurniawan ND, Reutens DC. A segmentation protocol and MRI atlas of the C57BL/6J mouse neocortex. Neuroimage. 2013;78:196–203. doi:10.1016/j.neuroimage.2013.04.008

36. Richards K, Watson C, Buckley RF, et al. Segmentation of the mouse hippocampal formation in magnetic resonance images. Neuroimage. 2011;58(3):732–740. doi:10.1016/j.neuroimage.2011.06.025

37. Manjón JV, Coupé P, Martí-Bonmatí L, Collins DL, Robles M. Adaptive non-local means denoising of MR images with spatially varying noise levels. J Magn Reson Imaging. 2010;31(1):192–203. doi:10.1002/jmri.22003

38. Zhao C, Tapera T, Cieslak M, Satterthwaite T. ModelArray: An R Package for Statistical Analysis of Fixel-Wise Data and Beyond.; 2025. https://pennlinc.github.io/ModelArray

39. Zhao C, Tapera TM, Bagautdinova J, et al. ModelArray: an R package for statistical analysis of fixel-wise data. Neuroimage. 2023;271:120037. doi:10.1016/j.neuroimage.2023.120037

40. R Core Team. R: A Language and Environment for Statistical Computing. R Foundation for Statistical Computing; 2024. https://www.R-project.org/

41. Dawson C. Ggprism: A “ggplot2” Extension Inspired by “GraphPad Prism.”; 2024. https://csdaw.github.io/ggprism/

42. Nilearn contributors, Chamma A, Frau-Pascual A, et al. nilearn. Published online June 21, 2025. doi:10.5281/ZENODO.15709579

43. Engler JB. Tidyplots empowers life scientists with easy code-based data visualization. iMeta. 2025;4(2):e70018. doi:10.1002/imt2.70018

44. Bethlehem RAI, Seidlitz J, White SR, et al. Brain charts for the human lifespan. Nature. 2022;604(7906):525–533. doi:10.1038/s41586-022-04554-y

45. Fujita S, Mori S, Onda K, et al. Characterization of Brain Volume Changes in Aging Individuals With Normal Cognition Using Serial Magnetic Resonance Imaging. JAMA Netw Open. 2023;6(6):e2318153. doi:10.1001/jamanetworkopen.2023.18153

46. Vinke EJ, De Groot M, Venkatraghavan V, et al. Trajectories of imaging markers in brain aging: the Rotterdam Study. Neurobiol Aging. 2018;71:32–40. doi:10.1016/j.neurobiolaging.2018.07.001

47. Ma Y. In vivo 3D digital atlas database of the adult C57BL/6J mouse brain by magnetic resonance microscopy. Front Neuroanat. 2008;2:175. doi:10.3389/neuro.05.001.2008

48. the Alzheimer’s Disease Neuroimaging Initiative (ADNI), EPIGEN Consortium, IMAGEN Consortium, et al. Identification of common variants associated with human hippocampal and intracranial volumes. Nat Genet. 2012;44(5):552–561. doi:10.1038/ng.2250

49. Heise V, Offer A, Whiteley W, Mackay CE, Armitage JM, Parish S. A comprehensive analysis of APOE genotype effects on human brain structure in the UK biobank. Transl Psychiatry. 2024;14(1):143. doi:10.1038/s41398-024-02848-5

50. Lyall DM, Cox SR, Lyall LM, et al. Association between APOE e4 and white matter hyperintensity volume, but not total brain volume or white matter integrity. Brain Imaging Behav. 2020;14(5):1468–1476. doi:10.1007/s11682-019-00069-9

51. Ruigrok ANV, Salimi-Khorshidi G, Lai MC, et al. A meta-analysis of sex differences in human brain structure. Neurosci Biobehav Rev. 2014;39:34–50. doi:10.1016/j.neubiorev.2013.12.004

52. Sang F, Chen Y, Chen K, Dang M, Gao S, Zhang Z. Sex differences in cortical morphometry and white matter microstructure during brain aging and their relationships to cognition. Cereb Cortex. 2021;31(11):5253–5262. doi:10.1093/cercor/bhab155

53. Sanchis-Segura C, Ibañez-Gual MV, Adrián-Ventura J, et al. Sex differences in gray matter volume: how many and how large are they really? Biol Sex Differ. 2019;10(1):32. doi:10.1186/s13293-019-0245-7

54. Guma E, Beauchamp A, Liu S, et al. Comparative neuroimaging of sex differences in human and mouse brain anatomy. Elife. 2024;13:RP92200. doi:10.7554/eLife.92200

55. Williams CM, Peyre H, Toro R, Ramus F. Neuroanatomical norms in the UK Biobank: The impact of allometric scaling, sex, and age. Hum Brain Mapp. 2021;42(14):4623–4642. doi:10.1002/hbm.25572

56. Clifford KP, Miles AE, Prevot TD, et al. Brain structure and working memory adaptations associated with maturation and aging in mice. Front Aging Neurosci. 2023;15. doi:10.3389/fnagi.2023.1195748

57. Young AL, Oxtoby NP, Daga P, et al. A data-driven model of biomarker changes in sporadic Alzheimer’s disease. Brain. 2014;137(9):2564–2577. doi:10.1093/brain/awu176

58. Young AL, Marinescu RV, Oxtoby NP, et al. Uncovering the heterogeneity and temporal complexity of neurodegenerative diseases with subtype and stage inference. Nat Commun. 2018;9(1):4273. doi:10.1038/s41467-018-05892-0

59. Archetti D, Young AL, Oxtoby NP, et al. Inter-Cohort Validation of SuStaIn Model for Alzheimer’s Disease. Front Big Data. 2021;4:661110. doi:10.3389/fdata.2021.661110

60. Poulakis K, Pereira JB, Muehlboeck JS, et al. Multi-cohort and longitudinal Bayesian clustering study of stage and subtype in Alzheimer’s disease. Nat Commun. 2022;13(1):4566. doi:10.1038/s41467-022-32202-6

61. Wheatley SH, Mohanty R, Poulakis K, et al. Divergent neurodegenerative patterns: Comparison of [18F] fluorodeoxyglucose-PET- and MRI-based Alzheimer’s disease subtypes. Brain Commun. 2024;6(6):fcae426. doi:10.1093/braincomms/fcae426

62. Lotze M, Domin M, Gerlach FH, et al. Novel findings from 2,838 adult brains on sex differences in gray matter brain volume. Sci Rep. 2019;9(1):1671. doi:10.1038/s41598-018-38239-2

63. Qiu LR, Fernandes DJ, Szulc-Lerch KU, et al. Mouse MRI shows brain areas relatively larger in males emerge before those larger in females. Nat Commun. 2018;9(1):2615. doi:10.1038/s41467-018-04921-2

64. Knickmeyer RC, Wang J, Zhu H, et al. Impact of sex and gonadal steroids on neonatal brain structure. Cereb Cortex. 2014;24(10):2721–2731. doi:10.1093/cercor/bht125

65. Qi G, Mi Y, Shi X, Gu H, Brinton RD, Yin F. ApoE4 impairs neuron-astrocyte coupling of fatty acid metabolism. Cell Rep. 2021;34(1):108572. doi:10.1016/j.celrep.2020.108572

66. Vitali F, Wiegand JP, Parker-Halstead L, Tucker A, Brinton RD. Weight trajectories in aging humanized APOE mice with translational validity to human alzheimer’s risk population: a retrospective analysis. PLOS One. 2025;20(1):e0314097. doi:10.1371/journal.pone.0314097

67. Kotredes KP, Pandey RS, Persohn S, et al. Characterizing molecular and synaptic signatures in mouse models of late-onset Alzheimer’s disease independent of amyloid and tau pathology. Alzheimers Dement. 2024;20(6):4126–4146. doi:10.1002/alz.13828

68. Reyes-Reyes EM, Brown J, Trial MD, et al. Vivaria housing conditions expose sex differences in brain oxidation, microglial activation, and immune system states in aged hAPOE4 mice. Exp Brain Res. 2024;242(3):543–557. doi:10.1007/s00221-023-06763-x

